# Anterior prefrontal cortex mediates implicit inferences

**DOI:** 10.1101/390427

**Authors:** Martijn E. Wokke, Tony Ro

## Abstract

Frequent experience with regularities in our environment allows us to use predictive information to guide our decision process. However, contingencies in our environment are not always explicitly present and sometimes need to be inferred. Heretofore, it remained unknown how predictive information guides decision-making when explicit knowledge is absent and how the brain shapes such implicit inferences. In the present experiment, participants performed a discrimination task in which a target stimulus was preceded by a predictive cue. Critically, participants had no explicit knowledge that some of the cues signaled an upcoming target, allowing us to investigate how implicit inferences emerge and guide decision-making. Despite unawareness of the cue-target contingencies, participants were able to use implicit information to improve performance. Concurrent EEG recordings demonstrate that implicit inferences rely upon interactions between internally and externally oriented networks, whereby anterior prefrontal regions inhibit right parietal cortex under internal implicit control.

**Significance:** Regularities in our environment can guide our behavior providing information about upcoming events. Interestingly, such predictive information does not need to be explicitly represented in order to effectively guide our decision process. Here, we show how the brain engages in such real-world ‘data mining’ and how implicit inferences emerge. We employed a contingency cueing task and demonstrate that implicit inferences influenced responses to subsequent targets despite a lack of awareness of cue-target contingencies. Further, we show that these implicit inferences emerge through interactions between internally- and externally-oriented neural networks. The current results highlight the importance of the anterior prefrontal cortex in transforming external events into predictive internalized models of the world.

## Introduction

Frequent exposure to regularities in our environment allows us to exploit consistencies to anticipate upcoming events. For instance, when strolling in an unfamiliar supermarket in search of a favorite chocolate bar, one typically does not pay much attention when passing by the detergents, but when the cookies come in sight, attention starts to focus. Without being explicitly told where to look for the product, the attentional system is able to use prior information (i.e., experience with supermarket layouts) and current sensory input to aid in the quest for chocolate. This example demonstrates that, in addition to externally observable information, internally oriented processes (e.g., memory, prospection) play a crucial role in efficiently guiding our behavior in everyday settings.

In order to understand decision-making in terms of network dynamics, it is essential to understand the mechanisms by which information is routed between brain regions. It has been proposed that alpha activity serves as a mechanism that gates the flow of information to relevant brain regions through inhibition (Vissers, 2018; Van Diepen et al., 2015; Matthewson et al., 2011; Matthewson et al., 2009; Jensen & Mazaheri, 2010; Klimesch et al., 2007; Fu et al., 2001). Alpha effects are typically measured after explicitly instructing participants about cues predicting a subsequent stimulus or indicating the location of an upcoming target (Foxe & Snyder, 2011; Worden et al., 2000), thereby mainly probing networks associated with external information processing (i.e., the dorsal attention network). In many cases, however, we learn to use predictive information in our environment in an implicit manner (Chun, 2000; Goldfarb et al., 2016), without the need of explicit knowledge about existing stimulus associations (Cleeremans et al., 1998; Cleeremans & Jiménez, 2002; Frensch & Rünger, 2003; Wokke et al., 2017). In such settings, internally oriented networks play an important role in formulating and testing of internally generated hypotheses (Christoff & Gabrieli, 2000) and comparing past and current sensory inputs (Wilson et al., 2014). To date it remains unclear how predictive information from our environment guides decision-making when explicit instructions are absent. Further, it is unknown how internally and externally oriented networks contribute to implicit inferences.

In the present study, we investigated how implicit contingencies guide decision-making. Participants performed an orientation discrimination task in which a target stimulus was preceded by a predictive cue. Critically, participants were not instructed and had no explicit knowledge that some of the cues signaled an upcoming target. Therefore, the information content of the cues was not ‘directly observable’ (Wilson et al, 2014; Schuck et al., 2016) and required information from previous trials (i.e., frequent exposure to cue-target pairings). During the task we recorded electroencephalographic (EEG) signals allowing us to measure whether implicit cues were able to influence behavioral responses, modulate alpha activity, and affect target processing, despite the fact that subjects were not explicitly aware of the meaning of the cues.

## Results

### Behavioral results

To determine whether participants were able to use implicit information to guide their behavior, we compared reaction time differences and differences in task performance (d’) between implicitly (validly) cued and implicitly (invalidly) non-cued targets (see Figure 1). We expected differences to occur specifically in the last half of the experiment when cue-target context had been established (i.e., after extensive exposure to pairings of target cue with the presentation of a target stimulus). Therefore, we split the data into the first and second halves of the experiment (Figure 2). Further, we assessed whether metacognitive performance (meta d’ and metacognitive efficiency) was affected by the implicit cues. For task performance there was a significant main effect of first/second half (block) of the experiment (*F*_(1,15)_= 8.75, *p*= 0.010). For both RT and task performance, there was a significant interaction effect between block and cue type (RT: *F*_(1,15)_= 5.17, *p*= 0.038; d’: *F*_(1,15)_= 14.43, *p*= 0.002). These interactions reflect differences in RT and performance that change over the course of the experiment depending on the cue type that preceded a target. To investigate these interactions further, we compared cued targets vs. non-cued targets for each half of the experiment separately using paired t-tests (two-tailed). As expected, there were no differences in the first half of the experiment for both RT (*t*_(15)_= 0.058, *p*= 0.955, BF_10_=0.256) and performance (*t*_(15)_= − 0.23, *p*= 0.821, BF_10_=0.262). In contrast, for the second half of the experiment, there were significant differences in RT and performance depending on cue type (RT: *t*_(15)_= − 3.144, *p*= 0.007, BF_10_=7.639; d’: *t*_(15)_= −3.058, *p*= 0.008, BF_10_=6.596; (Figure 2). These results demonstrate that participants learned to use the cues to increase the efficiency of their performance despite not having any explicit knowledge about the presence of cues. There was no difference in metacognitive performance between cued and non-cued targets (meta d’: *t*_(15)_= 1.041, *p*= 0.317; meta efficiency:: *t*_(15)_= 0.077, *p*= 0.940).

**Figure 1.**
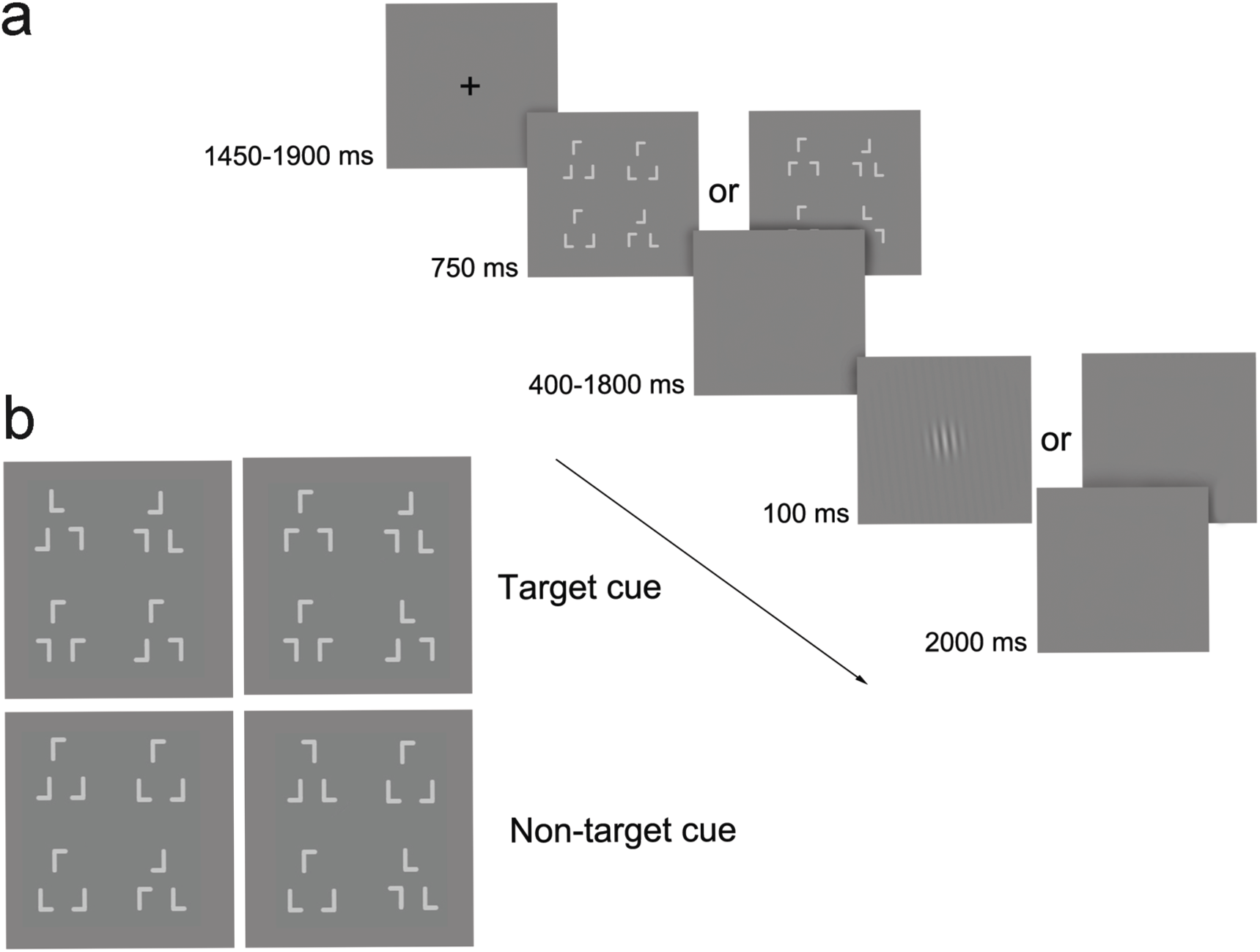
*a)* Participants had to respond as quickly as possible to a slightly right or left tilted vertical Gabor stimulus. Prior to target presentation a cue signaled either an upcoming target (100% validity) or a blank (66% validity). Participants were unaware of the relationship between the cue stimulus and target presentation during the experiment. *b)* Cues were made up of configurations of L-like shapes. The upper left and lower right configurations determined the identity of the cue (target or non-target cue).

**Figure 2.**
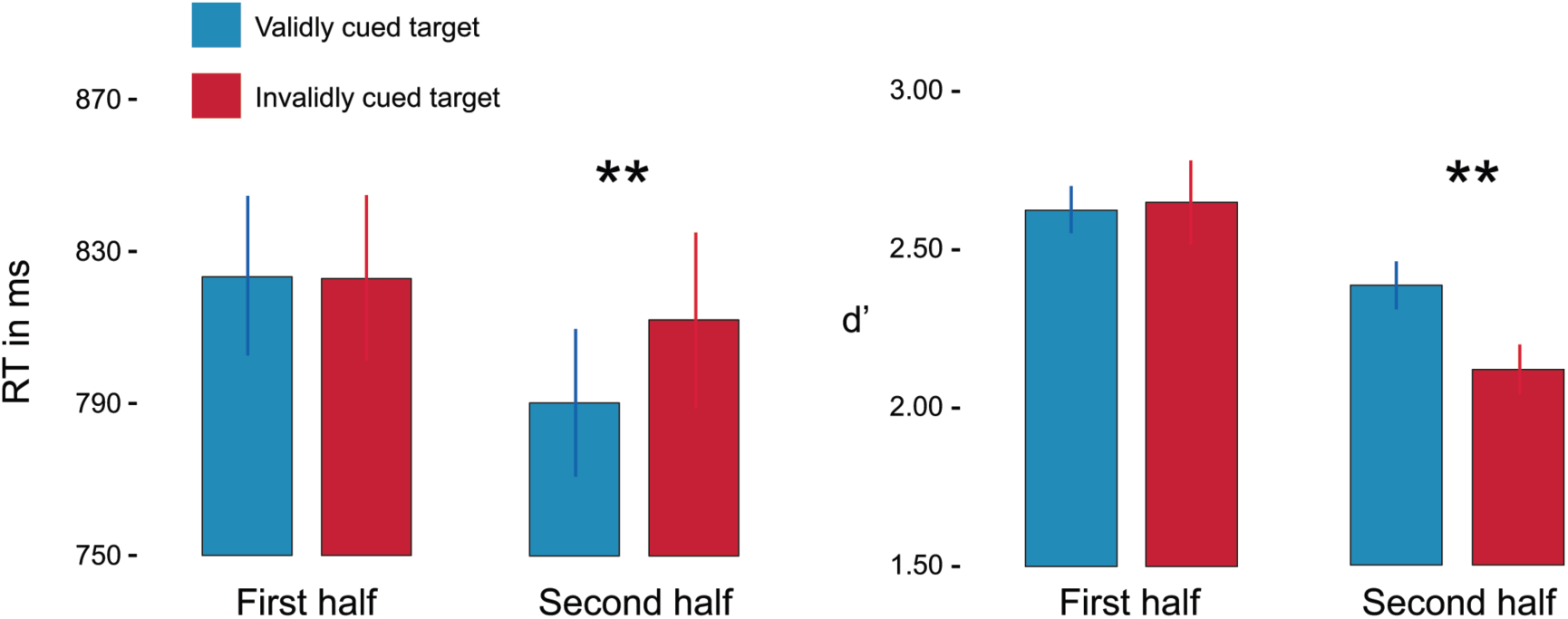
*a)* Participants responded faster (a) and performed better (b) when a target was preceded by a target cue than when preceded by a non-target cue. Bars are means +/- within subject SEM.

We repeated the same analyses for the second session, in which participants had explicit knowledge about the information conveyed by the cues. Importantly, we observed a significant main effect of cue for both RT (*F*_(1,15)_= 8.76, *p*= 0.010) and performance (*F*_(1,15)_= 12.57, *p*= 0.003). In addition, we also observed a block x cue interaction for RT (*F*_(1,15)_= 5.37, *p*= 0.035). Reaction times were only significantly faster for cued targets in the second half of the experiment (RT first half: *t*_(15)_= −1.74, *p*= 0.103, BF_10_=0.872; RT second half: *t*_(15)_= 3.57, *p*= 0.003, BF_10_=15.889), while performance was better in both the first and second halves of the experiment for cued targets compared to non-cued targets (d’ first half: *t*_(15)_= 2.50, *p*= 0.025, BF_10_=2.628; d’ second half: *t*_(15)_= 2.77, *p*= 0.014, BF_10_=4.052).

### EEG results

To determine whether alpha activity was influenced when implicit information guided behavior, we compared alpha power changes in a 400 ms time window after cue offset (before earliest target onset) between trials in which an implicit target cue and an implicit non-target cue was presented.

We observed a significant interaction (*F*_(4,60)_= 4.60, *p*= 0.003) between cue type (target/non-target) and channel location (O2, P4, C3, F4 and Fp2, see methods). These results demonstrate that depending on electrode location, there is a difference in alpha power between the two cues. There was lower alpha power over Fp2 (*t*_(15)_= −2.65, *p* =0.018, BF_10_=3.346) for target cues compared to non-target cues. In contrast, we observed a smaller alpha decrease in P4 for a target cue compared to a non-target cue (*t*_(15)_= 2.65, *p*=0.018, BF_10_=3.334; Figure 3b-c). When examining both cue types separately, we observed significant decreases of alpha power compared to baseline in P4 after both target and non-target cue presentation, while observing a significant alpha decrease in Fp2 exclusively when a target cue was presented (*t*s<-2.50, *p*s<0.025, BF_10_>2.73). For C3, O2, and F4, we observed no differences between cue types (all *t*s<0.972, all *p*s>0.346). These results demonstrate that although alpha decreased significantly after both a target cue and non-target cue in P4, there was less decrease of alpha in right parietal regions when a target cue was presented compared to non-target cue presentation. In contrast, there was a significant alpha power decrease in the right anterior frontal region exclusively after a target cue was presented (Figure 3c).

**Figure 3.**
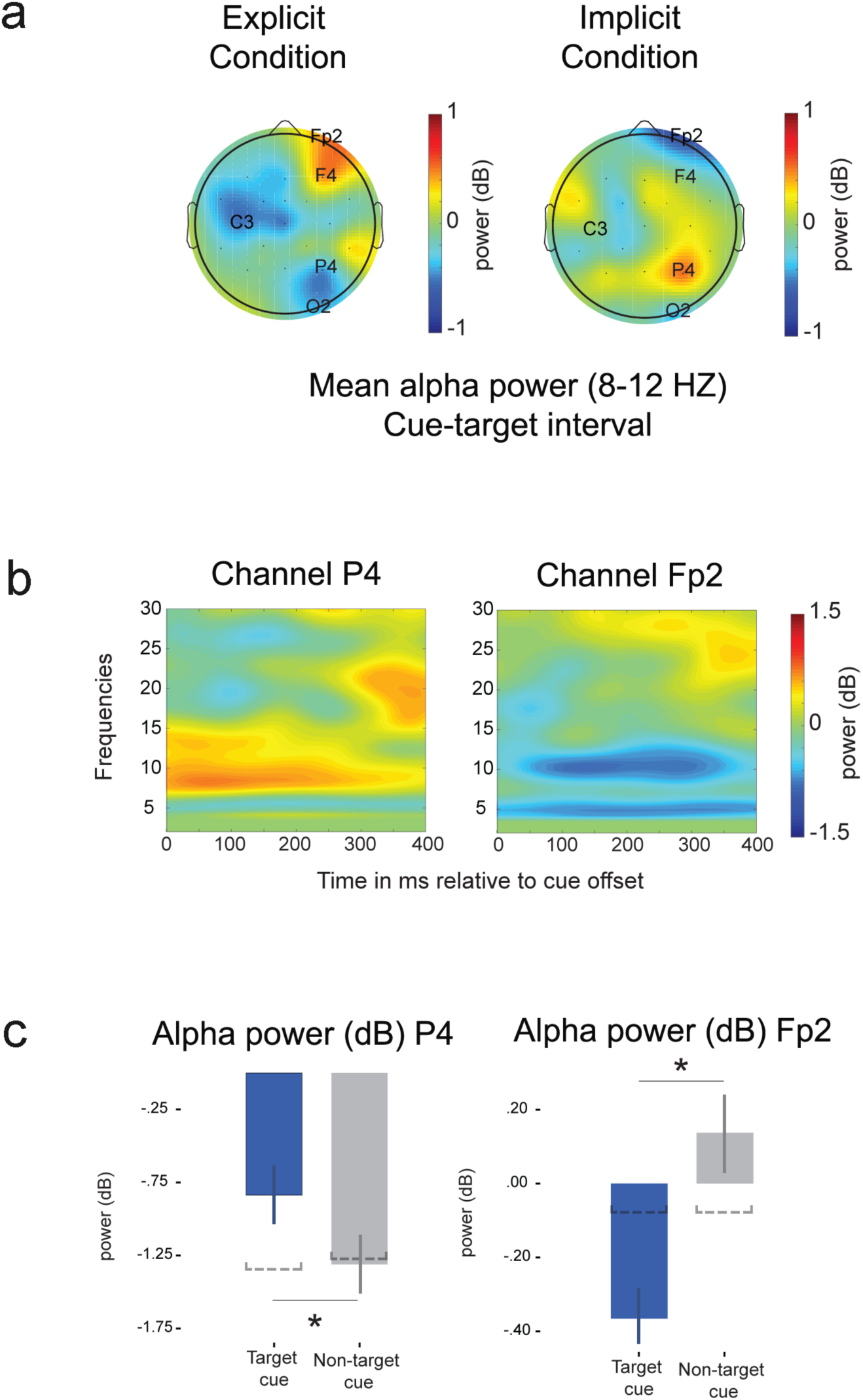
*a)* Electrodes with alpha activity differences between cue types in the explicit condition were used for further analyses (left). Topographic plot of alpha activity differences between cue types in the cue-target interval in the implicit condition (right). *b)* Time-frequency plot of electrodes P4 and Fp2 of differences between cue types. *c)* We observed a smaller alpha power decrease in right parietal region after target cue presentation compared to non-target cue presentation. In contrast, alpha power decreased in right orbitofrontal regions exclusively in response to a target cue. The dashed lines represent the mean values observed in the first half of the experiment. Bars represent means +/- within subject SEM.

To examine this effect further and test whether the observed cue type effects in P4 and Fp2 related to implicit learning of cue-target contingencies and the build-up of task context, we investigated whether alpha power differences between cue types could be observed in the first half of the experiment. No differences were observed in either P4 or Fp2 between the two cue types (*t*s<0.339, all *p*s>0.744), see Figure 3c (dashed lines). For both cue types we observed decreased alpha compared to baseline in P4 (target cue: *t*_(15)_= −3.31, *p* =0.005, BF_10_=16.296; non-target cue: *t*_(15)_= −3.59, *p* =0.003, BF_10_=10.097). However, no alpha changes were found in Fp2 for both cue types (*t*s<0.53, all *p*s>0.607).

Next, we explored data from the explicit cueing condition to further scrutinize the alpha power effects we found in the implicit condition. In the explicit condition, attention could be externally oriented. Therefore, we expected to find typical alpha power effects (i.e., an alpha power decrease in P4 after target cueing, see Heilman & Van Den Abell, 1980; Sacchet et al., 2015; Zago et al., 2016; Lobier et al., 2018) in the explicit cueing condition. We investigated alpha activity changes (compared to baseline) in the explicit condition only in P4 and Fp2 for both target and non-target cues. We observed a significant decrease of alpha power compared to baseline in P4 exclusively after target cue presentation (*t*_(15)_ = −4.01, *p* =0.001, BF_10_=33.236; for non-target cue: *t*_(15)_= −2.00, *p*=0.063, BF_10_=1.335). No significant alpha changes were observed in Fp2 for both cue types (*t*s<1.68, *p*s>0.113, BF_10_<0.881).

To examine interactions between different cortical regions, we assessed measures of interregional functional connectivity (alpha phase synchrony) by calculating intersite phase clustering (ISPC) between ‘seed’ P4 and O2, C3, F4 and Fp2 (see methods). There was a significant main effect of channel location (*F*_(4,45)_= 4.89, *p*= 0.005) and a marginally significant interaction between cue type and channel location (*F*_(4,45)_= 2.39, *p*= 0.081). We observed increased alpha-band synchronization between P4 and Fp2 (*t*_(15)_= 3.08, *p*=0.008, BF_10_=6.892) after a target cue was presented in comparison to a non-target cue, see Figure 4 (we plotted ISPC differences between P4 and all other electrodes for illustration purposes, while only statistically testing ISPC changes between P4 and O2, C3, F4, and FP2). We observed no significant differences between P4 and the other electrodes (all *t*s<1.31, all *p*s>0.208).

**Figure 4.**
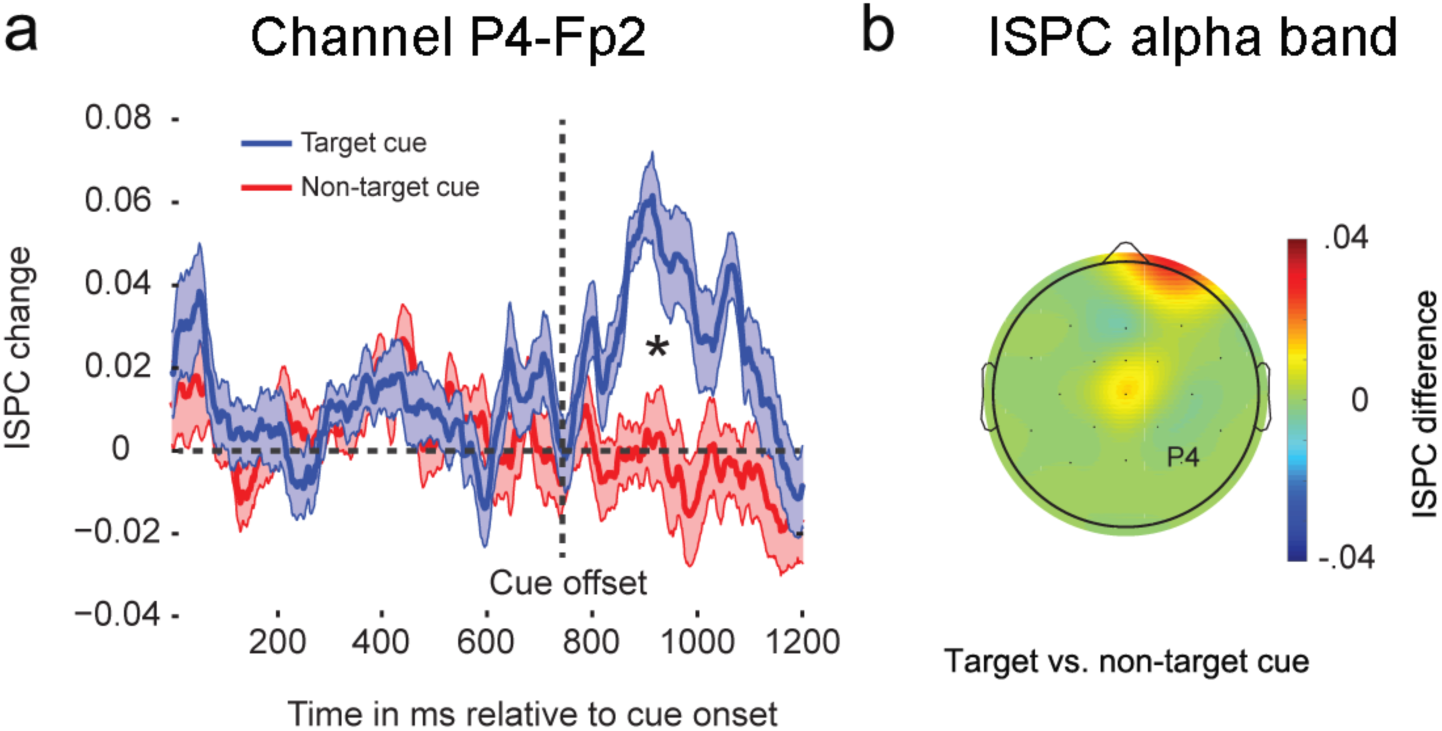
*a)* We observed significantly larger alpha phase synchrony between P4 and Fp2 for target cues in the cue-target interval. We plotted the period from cue onset to target onset for illustration purposes, while only comparing mean ISPC changes in the interval after cue offset (shaded areas are +/− within subject SEM). *b)* Topographic plot of mean ISPC changes in the cue-target interval (target vs. non-target cue) using P4 as the ‘seed’ electrode.

To determine how neural measures relate to behavior, we correlated RT and d ‘ differences to cued and non-cued targets with alpha power decreases after target cue offset for P4 and Fp2 and parietal-anterior frontal functional connectivity changes after target cue presentation. We observed a significant correlation between parietal-anterior frontal ISPC change and the RT effect (*r*= 0.769, *n*=16, *R^2^*= 0.59, FDR<0.05, BF_10_=78.73; see Figure 5). We did not find any significant correlations that survived the multiple comparisons correction between RTs and alpha power changes or between d’ and ISPC change (all *r*s<0.335, FDR>0.05). These findings demonstrate a strong link between enhanced alpha phase synchrony between P4 and FP2 and the speeding of responses due to implicitly learning of cues predicting an upcoming target stimulus.

**Figure 5.**
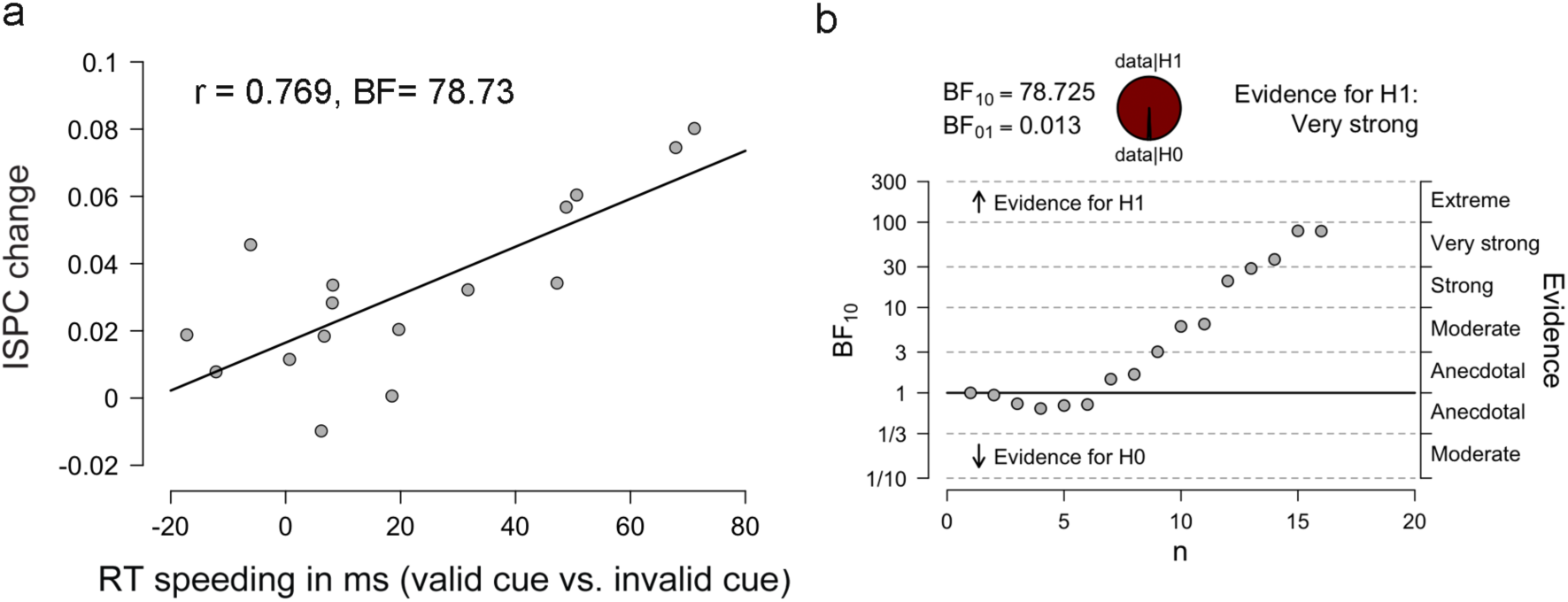
**a)** RT decreases are highly correlated with functional connectivity changes between P4 and Fp2. **b)** Sequential analysis of the Bayesian correlation pairs illustrates the strength of the effect and the amount of participants included.

Finally, we investigated whether neural signals related to target processing differentiated depending on preceding cue type (target vs. non-target cue) in both the implicit and explicit conditions (see methods). We examined whether we could find differences in P3a and P3b ERP components associated with ‘context updating’ of the stimulus environment (Donchin, 1981; Polich & Kok, 1995), and linked to differences in access awareness (Naccache et al., 2016). We observed a significant cue (target/non-target cue) x ERP type (P3a/P3b) x session (implicit/explicit) interaction (*F*_(1,13)_= 8.95, *p*= 0.010). Interestingly, in the implicit condition we found an increased P3a when a target was preceded by a non-target cue compared to when a target was preceded by a target cue (*t*_(13)_= 3.61, *p*=0.003, BF_10_=14.627), see Figure 6a. In contrast, we found an increased P3b in the explicit condition when a target was preceded by a non-target cue compared to when a target was preceded by a target cue (*t*_(13)_= 3.22, *p*=0.007, BF_10_=7.902), see Figure 6d. These results seem to corroborate previous findings demonstrating the influence of contextual processes on the P3, where P3 activity is modulated when the model or context of a stimulus environment needs to be updated (Donchin, 1981; Donchin and Coles, 1988; Polich & Kok, 1995; Silverstein et al., 2015; Todorovic et al., 2011; Seppänen et al., 2012; Bang & Rahnev, 2017; Diaz-Brage et al., 2018).

**Figure 6.**
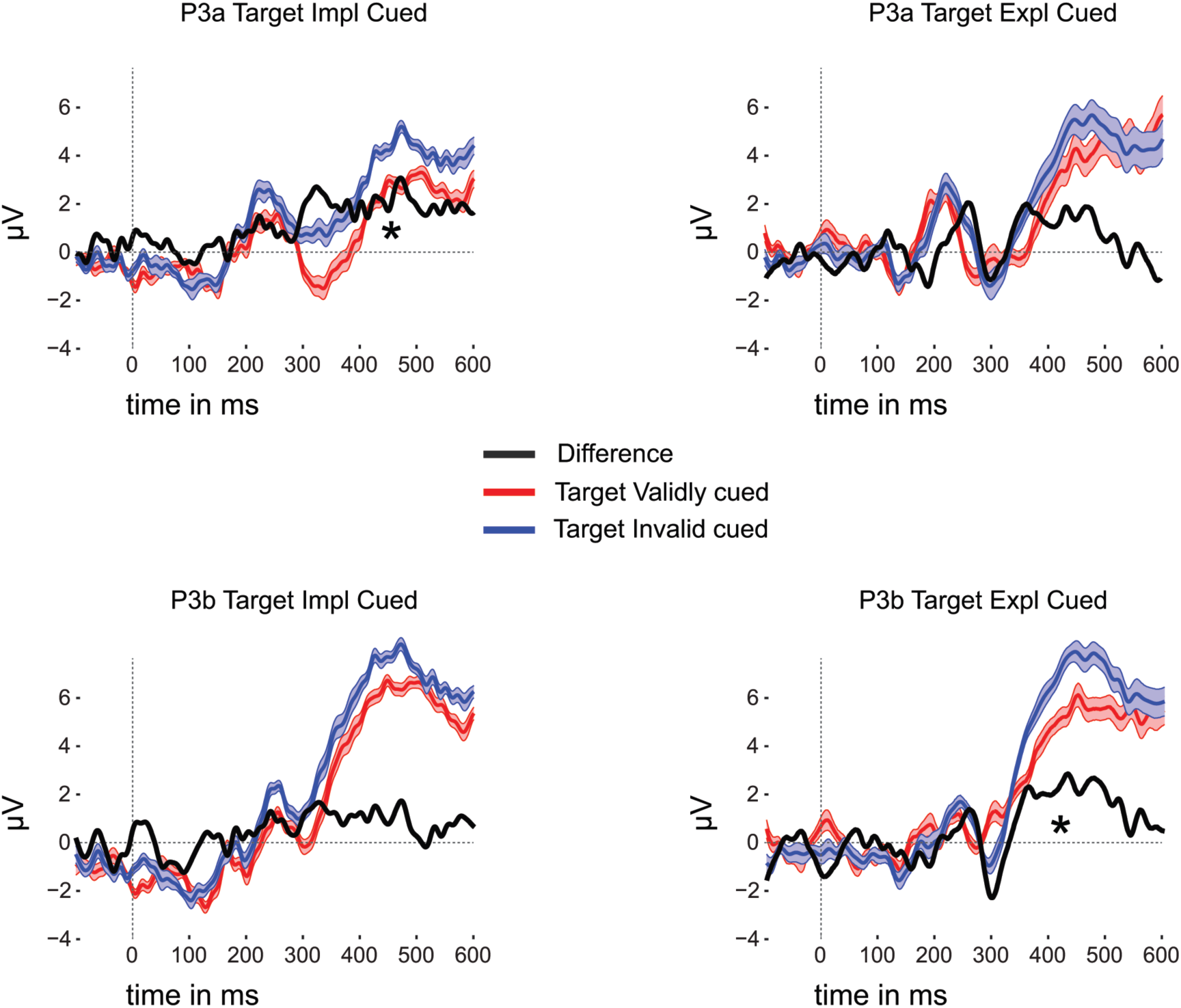
ERPs to targets on trials preceded by a target cue resulted in a smaller P3a in the implicit condition (a) and a smaller P3b in the explicit condition (d). Shaded areas are +/− within subject SEM.

## Discussion

In everyday life, we are able to use predictive information in our environment to guide our behavior. However, sometimes information is not readily available and needs to be inferred (O’Doherty et al., 2001; Wilson et al., 2014). In such cases, it is necessary to compare past and current sensory inputs and use prior experience to select relevant information to anticipate upcoming events (Chun et al., 2011; Wilson & Niv, 2012).

In this experiment, implicit cues were used to investigate how unconscious contingencies may be able to control our decision process. Specifically, we focused on whether implicit cueing was able to affect behavioral responses in a discrimination task and modulate oscillatory neural activity in the alpha frequency range. Results demonstrate that participants were able to use implicit cues to improve performance and speed up responses (Figure 2; Chang et al., 2015; Pinto et al., 2015; Stein & Peelen, 2015; Meijs et al., 2018). We observed a specific decrease of right anterior frontal alpha power when a target stimulus was implicitly cued (Figure 3c), whereas alpha power decrease over right parietal cortex diminished after the presentation of an implicit target cue. These findings corroborate previous findings demonstrating that anterior frontal cortex becomes recruited when information needs to be inferred (Wilson et al., 2014; Shucks et al., 2017; Christoff & Gabrieli, 2000). Furthermore, it has been shown that alpha power increases in parietal cortex when attention becomes internally oriented (Ray & Cole, 1985; Schupp, 1994; Cooper et al., 2003). Interestingly, we observed a specific increase in functional connectivity (alpha phase synchrony) between right parietal and right anterior frontal regions when implicit information was used (Figure 4). This change in functional connectivity in response to an implicit target cue correlated strongly with behavioral effects (Figure 5). Finally, ERP differences (Figure 6) between cued and non-cued targets showed that cued targets were implicitly anticipated (Summerfield et al., 2008; Todorovic et al., 2011; Chennu et al., 2013). Figure 7 summarizes these results and provides a schematic of the mechanisms mediating implicit inferences.

**Figure 7.**
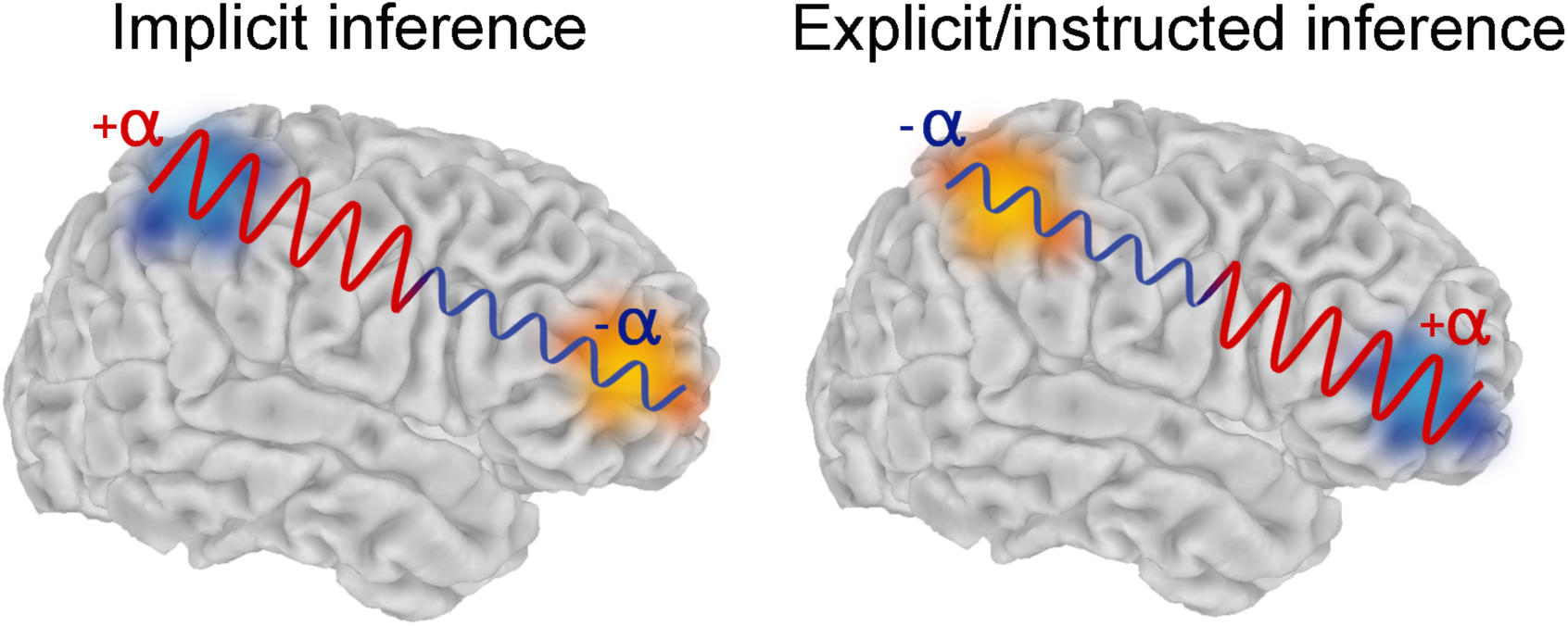
Implicit inferences (top) engage internally oriented networks, enhancing processing via anterior prefrontal regions. Competitive network dynamics result in decreased externally oriented network activity, where alpha activity serves as a mechanism to gate the flow of information within specific networks. Explicit instructed inference (bottom) results in an opposite pattern, whereby externally oriented networks are engaged.

### Alpha oscillations and gating

Alpha activity has long been considered a marker for increased inhibition (Lopes Da Silva, 1991). Recently, it has been put forward that alpha oscillations play a key role in the gating of the flow of information by suppressing processing of information in task-irrelevant networks (Mathewson et al., 2014; Jensen & Mazehari, 2010; Klimesch et al., 2007). It has been demonstrated that a shift of attention to either the left or right visual hemifield decreases alpha in the contralateral hemisphere, while increasing alpha in the ipsilateral hemisphere (Sauseng et al., 2005; Thut et al., 2006; Worden et al., 2000). Furthermore, recent studies have shown that alpha power increases in the dorsal stream when a task relies on ventral stream processing (Jokish and Jensen, 2007; Wokke et al., 2014). In the present study, we observed a smaller decrease of parietal alpha after presentation of a target cue when participants became sensitive to implicit cueing (Figure 3). It has been previously shown that alpha in parietal regions increases when attention becomes internally directed, suggesting the necessity of active inhibition of external sensory input for internally driven mental operations (Ray & Cole, 1985; Schupp, 1994; Cooper et al., 2003). Further, Sestieri et al. (2010) observed functional competition between internally (memory) and externally (perception) driven processes, where parietal cortex operated in a push-pull manner depending on the task engaging either internally (search in memory) or externally (search in the environment) oriented networks. Here, enhanced internally driven processes could dampen typical parietal alpha power decreases due to functional competition (Fox et al., 2005). This push-pull hypothesis seems to be supported by the recruitment of anterior frontal regions, strongly associated with the evaluation of internally generated information (Shucks et al., 2017; Christoff & Gabrieli, 2000; Dixon et al., 2018). Further, a seemingly opposite pattern was observed in the explicit condition, when attention could be externally oriented (see Figure 3a and Figure 7).

Lesions to orbitofrontal cortex have been shown to induce a state in which subjects become solemnly dependent on information from the outside world that is directly observable (utilization behavior, see Lhermitte, 1983; Brazzeli et al., 1998; Besnard et al., 2010), while internal models and information about context are no longer accessible (similar effects have also been observed in reversal learning tasks, see Dias et al, 1998 and Wilson et al., 2014). These findings have been associated with a disrupted balance in network functioning, where orbitofrontal damage results in a disinhibited state of parietal cortex (Lhermitte et al., 1986). Interestingly, the current results show increased functional connectivity between parietal and orbitofrontal regions, exclusively in the implicit condition (Figures 3 & 4). Measures of functional connectivity provide us with information about formation and functional integration of networks, working either in concert or in a push-pull fashion (Wokke et al., 2015; Srinivasan et al., 2007; Fox et al., 2005). In the last decades, competing network dynamics have been demonstrated by opposed activity levels in intrinsic ‘outward oriented’ networks and the ‘internally oriented’ default-mode network on a variety of tasks (Raichle et al., 2001; Fox et al., 2005; Weissman et al., 2006; Kelly et al., 2008; Hampson et al., 2010; Wokke et al., 2015). As depicted in Figure 7, our results indicate a similar dynamic of internally and outward oriented network activity when participants learn to use implicit information. Further, the strength of the functional connectivity change between parietal and anterior/orbitofrontal cortex strongly correlated with the behavioral RT effect. However, we did not find a significant correlation between the behavioral effect and alpha power in either parietal or anterior/orbitofrontal regions. These findings indicate that specifically the orchestration of activity in internally (anterior frontal) and externally (parietal) oriented networks could be fundamental for situations when information needs to be inferred.

### Orbitofrontal cortex and inferential decision-making

Activity in the orbitofrontal cortex (Brodmann areas 10, 11 and 47) has been consistently associated with support of adaptive decision-making by uncovering predictive values associated with stimuli in our environment (Walton et al., 2010; Boorman et al., 2016). The connectivity between orbitofrontal cortex and sensory, frontal, striatal, and hippocampal regions makes this region highly suited for the generation and testing of hypotheses (Frey & Petrides et al., 2002; Bar et al., 2006) and for providing predictions about specific outcomes associated with stimuli (Rudebeck & Murray, 2014; Goldfard et al., 2016). Recently, the above-described observations have been captured in a ‘state-space’ model in which the orbitofrontal cortex plays a crucial role. This state-space theory of orbitofrontal cortex (Gershman & Niv, 2010; Wilson et al., 2014; Schuck et al., 2016; Schuck et al., 2017) focuses on the context in which decisions are being made and what the decision-making agent considers ‘the state of the world’ at the moment of the decision (Schuck et al., 2017). Such states can be connected to external information (e.g., explicit cues) or they can contain internally generated information, which cannot be directly obtained from the immediate environment and has to be inferred (e.g., implicit cues or task context). Specifically, the orbitofrontal cortex seems critical for the representation of states that include such partially observable information (Brown et al., 2010; Wilson et al., 2014). Our current results suggests that anterior frontal cortex becomes recruited when ‘hidden’ information is available that can help optimize the decision process. The present findings are contributing to a growing amount of evidence demonstrating the critical role orbitofrontal cortex plays in using information in the environment that is not directly observable.

### P3a/P3b: Prediction and access consciousness

To further examine the consequences of implicit cueing, we investigated how cued and non-cued target stimuli influenced P3a/P3b activity. A rich literature describes the role of the P3 in context updating, (Donchin, 1981), in which a current stimulus is compared with a preceding stimulus in working memory (Donchin, 1981; Donchin and Coles, 1988; Polich & Kok, 1995; Silverstein et al., 2015). We therefore compared P3a and P3b responses to cued and non-cued targets. These P3 components have been frequently studied using ‘oddball’ designs, linking the P3 to updating of stimulus context (Donchin, 1981; Summerfield et al., 2008; Todorovic et al., 2011; Chennu et al., 2013). Interestingly, we observed an enhanced P3a when a target stimulus was preceded by a non-target cue in the implicit condition, whereas we found an increased P3b in the explicit condition.

It has been suggested that the P3a component relies more on automatic (unconscious) processes (Muller-Gas et al., 2007), whereas the P3b component is linked to access consciousness (Faugeras et al., 2012; Naccache et al., 2016, but see Silverstein et al., 2015). These findings are in line with a recent study investigating the relationship between top-down expectations and access consciousness (Meijs et al., 2018). In that study, the authors observed that access awareness of a predictive stimulus is necessary to actively use top-down predictions for subsequent target processing (in an attentional blink design where T1 predicted T2). The present results and the findings from Meijs et al. (2018) indicate that a predictive stimulus needs to be *perceptually* processed all the way up to the level of access awareness to be effective, but that the meaning of the stimulus (i.e., that the stimulus is in fact predictive) can still remain inaccessible for introspection without discarding its functionality. Further, Meijs et al. (2018) demonstrated that prediction errors could be triggered outside of conscious awareness. In the current study, we observed related effects by observing a P3a difference between cued and non-cued targets in the implicit condition, while we found a P3b difference in the explicit condition that was not present in the implicit condition. These findings suggest that unconscious/implicit ‘context updating’ effects proceed more automatically than in its conscious/explicit form (Faugeras et al., 2012; Naccache et al., 2016).

Previous work suggests that unconscious/automatic elicited responses are relatively short-lived while conscious detection results in more long-term behavioral adaptations (Cohen et al., 2009; Van Gaal et al., 2012), although it remains debated what the consequences of such differences exactly are. It would be interesting to investigate how long-lived the observed effects of implicit learning are (e.g., by testing participants on multiple occasions in the implicit condition to examine the longevity of the effect of implicit learning). In the present study, we also did not focus on how or when implicit control of attention became accessible for introspection. It would be very fascinating to investigate how the use of implicit information progresses towards explicit knowledge, and observe whether such a transition would proceed in a gradual or in an all-or-none manner (Sergent et al., 2004; Windey et al., 2015; King et al., 2016). It could be that hypotheses about implicit information gradually become strong enough, reaching increasingly higher signal-to-noise levels, resulting in stable (neural) representations (Schurger et al., 2010) and updating of internal predictive models of the environment (O’Reilly et al., 2013). Such internalization of stimulus-outcome events (Buzsáki et al., 2014; Wokke et al., 2017; Cleeremans, 2011) could pave the way for implicit information to become accessible for introspection.

## Conclusion

In daily life, our decisions are frequently guided by regularities in our environment. However, such contingencies are not always explicitly present and sometimes need to be inferred. Using contingency cueing, we show that implicit inferences influenced responses to subsequent targets despite a lack of awareness of cue-target contingencies. These implicit inferences emerge through changes in internally- and externally-oriented neural networks. The current results demonstrate that anterior prefrontal cortex plays an important role in the transformation of externally driven stimulus-outcome events into predictive internalized models of the world.

## Materials and Methods

### Participants

Seventeen participants (9 females, mean age= 25.4, SD= 6.3) took part in this study for financial compensations. All participants had normal or corrected-to-normal vision and were naïve to the purpose of the experiment. All procedures complied with international and institutional guidelines and were approved by the Institutional Review Board of The City University of New York. Prior to the experiment, participants were instructed on the task, after which all participants provided their written informed consent.

### Task design

Stimuli were presented full screen (1024 × 768 pixels) on a 17-in. CRT monitor (Sony Trinitron Multiscan 220GS) with a refresh rate set at 100 Hz. The monitor was placed at a distance of ~57 cm in front of each participant. Each trial started with a centrally presented fixation cross that was presented for 1455, 1685, or 1915 ms, after which the cue was presented for 750 ms. The cue consisted of four configurations of four L-shaped figures (Figure 1b) presented in each quadrant of the screen. After presentation of the cue, a blank screen was presented for 400, 800, 1200 or 1600 ms, after which a target or another blank screen was presented for 100 ms. Participants were instructed to keep their eyes open and to minimize blinks from cue onset until they gave their response to the target or the end of the trial (see Figure 1a). A target stimulus consisted of a slightly left or right tilted vertical Gabor patch (see Figure 1). We tilted the Gabor between 1-3^°^ to ensure that performance was kept below ceiling and above chance (~80% correct during practice trials, see below). After target presentation, participants had to indicate as quickly as possible the orientation of the Gabor (left or right) by pressing a corresponding left or right response button. Next, participants provided their confidence about their decision, on a scale ranging from 1 to 4 by pressing one of four buttons. Participants were instructed to assign a low value to a decision that was accompanied by low confidence in being correct and a high value when they were very confident about being correct. Participants were encouraged to make use of the whole scale. On trials when no target was presented, the participants were instructed not to respond and wait for the onset of the next trial (2 sec). We customized two (computer) mice in order to create four response buttons that registered responses through a Teensy LC board at microsecond temporal resolution.

A target cue (Figure 1b) always predicted an upcoming target (100% valid), whereas a non-target cue was followed by a target on one-third of the trials or no target on two-thirds of the trials). The upper right and lower left configuration of the cues determined cue type. Participants performed two separate sessions at least/approximately one week apart. Crucially, in the first session participants were not instructed about the types of cues signaling target stimuli or about the general purpose of the cue stimuli in the experiment. The cue parameters were based on data from a pilot study (n=60, spread over 5 different sessions) and were set such that participants were able to learn the contingencies between cue and target without gaining explicit knowledge about the meaning of the cue stimuli (i.e., explicitly recognize them as being cues). In the second session we explicitly instructed participants about the identity of the cues, explaining to the participants that the upper right and lower left configuration of each cue was predictive of trial type and which cue was most likely to be followed by a blank.

In both sessions, participants started with 120 trials of practice to get accustomed to the task. At the end of the first session, we determined whether participants gained explicit knowledge about the nature of the cue stimuli. In four steps we probed participants’ knowledge about the cues. First, we asked participants whether they noticed anything about the stimuli appearing in the experiment. Second, we asked whether they noticed if the stimulus with the L shaped figures had any purpose in the experiment. Next, we asked whether they noticed if specific configurations of L shapes signaled an upcoming target or whether configurations of specific L shapes were more related to the appearance of a blank. Finally, we showed participants the cues and tested if they could tell the difference between the cues and their relation to target presentation. Of all seventeen participants, only one noticed a relationship between the cues and the appearance of a target stimulus. For the other sixteen participants, there was no explicit knowledge of the presence of cues on any of the above-described questions (cf. Chun & Jiang, 2003; Geyer et al., 2012; Goujon et al., 2013). All analyses were based on these sixteen participants.

In each session, we presented 720 trials equally divided over 6 blocks. Within each block, 48 (validly) cued target trials, 24 (invalidly) non-cued target trials, and 48 (validly) cued blank trials were presented in pseudo-random order. Participants took a 10-minute break after completing three blocks. Each session lasted approximately two hours.

### Behavioral analyses

To assess whether response times, target discrimination accuracy, and metacognitive judgments differed depending on implicit cue type, we calculated reaction times, task performance (d’, see Macmillan & Creelman, 2004), metacognitive sensitivity (meta-d’), and metacognitive efficiency (meta-d’ – d’, Maniscalco & Lau, 2012; Fleming & Lau, 2014). Because first-order task performance is known to influence metacognitive sensitivity (Fleming & Lau, 2014), it is necessary to assess metacognitive sensitivity relative to different levels of first-order task performance (metacognitive efficiency). We performed three seperate 2 (first and second half of the experiment) x 2 (target and non-target cue) repeated measures ANOVAs on reaction times, performance (d’), and second-order task performance (meta-d’). Unfortunately, confidence judgments of two participants were not registered due to a technical error, basing the second-order performance analyses on 14 participants. All behavioral analyses were performed using Matlab (Matlab 12.1, The MathWorks Inc.), type 2 SDT scripts (Maniscalco & Lau, 2012) and SPSS (IBM SPSS Statistics, 22.0).

### EEG measurements and analyses

EEG was recorded and sampled at 1000 Hz using a 32-channel Easy Cap system (Easy Cap – Munich). Two additional electrodes were placed on the outer eye canthi to record eye blinks. Electrode impedance was kept below 20 kΩ. Offline, the data was high-pass (0.5 HZ) and low-pass (40 HZ) filtered and then re-referenced to the left and right mastoid. The data was epoched −0.7 to + 1.7 sec around cue onset. These time windows avoided edge artifacts resulting from time–frequency decomposition (see below). We removed trials containing irregularities due to eye blinks or other artifacts by visually inspecting all trials. To increase spatial specificity and to filter out deep sources we converted the data to spline Laplacian signals (Perrin et al., 1989; Cohen 2015).

As we expected to measure implicit contingency effects in the second half of the experiment, the last 70 target cue with target and last 70 non-target cue without target trials were selected after artifact rejection for all analyses. We decomposed the cue-locked, epoched EEG time series for these trials into their time–frequency representations by convolving them with a set of Morlet wavelets (frequencies ranging from 1 to 30 Hz in 1 Hz steps). Complex wavelets were created by multiplying perfect sine waves with a Gaussian. The range of the width of the Gaussian was set between 4 and 10 in 40 logarithmically scaled steps, in order to have a good trade-off between temporal and frequency resolution for each frequency. We applied the fast Fourier transform to the EEG data and the Morlet wavelets, after which these were multiplied in the frequency domain. Next, the inverse FFT was applied, allowing us to define an estimate of frequency-specific power at each time point and an estimate of the frequency-specific phase at each time from the resulting complex signal (Van Driel et al., 2015). We normalized the data (dB Power tf = 10 ^*^ Log10[Power tf / Baseline Power f]) using an interval of –300 to 0 ms relative to cue onset as baseline. For our hypothesis, we specifically focused on signals in the alpha frequency band between cue offset and earliest target onset (i.e., a time window of 0-400 ms after cue offset). To limit the amount of comparisons we preselected electrodes for our analyses in the implicit condition based on an independent dataset (EEG data from the explicit condition). Electrodes O2, P4, C3, F4 and Fp2 were selected based on differences in alpha power between trials containing a target cue vs. trials containing a non-target cue in the explicit condition (Figure 3a).

To further examine the way information might be gated via alpha oscillatory mechanisms, we assessed measures of interregional functional connectivity in the alpha range. Consistencies of the difference of time–frequency phase values between two channels in the alpha band across trials were computed (Intersite Phase Clustering (ISPC), see Siegel et al., 2012; Cohen, 2014). We chose P4 as our ‘seed’ electrode based on previous studies demonstrating the involvement of right parietal cortex in attention and alpha oscillations (Behrman et al., 2004; Bareham et al., 2018; Thut et al., 2011). We used the same preprocessing steps as described above for the time-frequency analyses and a baseline period of −300-0 ms before cue onset for both cue types.

We performed a 5 (channel location) x 2 (target and non-target cue) repeated measures ANOVA on mean alpha power changes and a 4 (channel location) x 2 (target and non-target cue) repeated measures ANOVA on mean ISPC changes.

Finally, we were interested whether implicit information influenced neural signals related to target processing. Therefore, we epoched the EEG data from −100 to 600 ms around target onset, using the same preprocessing steps as described above. Unfortunately, for two participants there were too many artifacts (> 50% of trials) in the epoch after target presentation, likely because of the long interval between cue onset and response in which we instructed participants not to blink, resulting in fourteen participants for our target ERP analyses. We focused on the P3a and P3b component, which have been shown to be highly associated with stimulus environment updating processes (i.e., comparing present and previous stimuli in working memory) and differences in levels of access consciousness, respectively (Donchin, 1981; Polich & Kok, 1995; Sergent et al., 2005; Muller-Gas et al., 2007; Naccache et al., 2016; Wokke et al., 2016). In light of findings demonstrating that the P3b indexes different levels of access consciousness, we tested ERP differences in both the implicit as well as the explicit condition. For the P3a component we selected Cz between 250-400 ms, while we selected Pz between 350-500 ms after target onset for P3b comparison (Polich, 2007).

All signal-processing steps were completed using EEGlab (Delorme & Makeig, 2004) and **X** code (Cohen, 2014) in Matlab (Matlab 12.1, The MathWorks Inc.), and statistical analyses were performed using Matlab (Matlab 12.1, The MathWorks Inc.), JASP (Version 0.8.6) and SPSS (IBM SPSS, Version 20.0).

